# Phenotypic selection analysis and confounding environmental variables

**DOI:** 10.1101/2022.06.15.496257

**Authors:** Michael B. Morrissey, Jonathan M. Henshaw

## Abstract

When environmental variation contributes to relationships between traits and fitness, it can confound analyses of phenotypic selection and, ultimately, bias predictions of adaptive evolution. To date, discussions of how to combat this problem emphasise complex statistical analyses aimed at estimating the genetic basis of the relationship between traits and fitness (e.g., the secondary theorem of selection). This article presents a path analysis-based description of the environmental confounding problem, which clarifies the issue and motivates simpler analyses as potential solutions. We show how standard selection analyses can be expanded to explicitly include environmental variables that may confound trait-fitness relationships, potentially leading to dramatically improved predictions of the evolutionary response to selection relative to classical phenotype-based estimates. We provide both univariate and multivariate treatments of the decomposition of the selection differential into components that may cause evolution via direct and indirect selection and components representing evolutionarily inert, environmentally-induced covariance. The multivariate treatment also yields expressions for the decomposition of the selection differential based on extended selection gradients, which may be of wide general interest beyond the environmental problem. Our approach to the environmental confounding problem makes more plausible demands on data than previous, more involved, quantitative genetic approaches, and addresses the issue of environmental confounding in a more biologically informative way.

## Introduction

Lande and Arnold (1983) presented a simple and transparent method for estimating selection gradients via the multiple regression of relative fitness on trait values. The method has since been used extensively to estimate the strength and direction of directional selection in nature (Kingsolver *et al*. 2001, Siepielski *et al*. 2009, 2017). A key justification – both for Lande’s (1979) definition of the direct selection gradient, *β*, and for Lande and Arnold’s (1983) multiple regression approach to estimating such gradients – is a simple equation that translates selection gradients into quantitative predictions about evolutionary change. This expression, known as the ‘Lande equation’, is

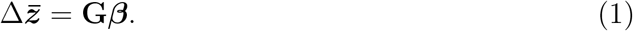

Here, per-generation evolutionary change of the mean phenotype, 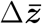, is given by the (matrix) product of the genetic variances and covariances of the traits in question, **G**, and the vector of selection gradients, ***β***. Importantly, Lande and Arnold (1983) provided a simple and remarkably robust means of estimating selection gradients: ***β*** is estimated as the coefficients from the multiple regression of relative fitness on trait values. This approach has been very widely used, yielding thousands of estimates of selection gradients for different traits (e.g., as synthesised in large meta-analyses such as Kingsolver *et al*. 2001, Siepielski *et al*. 2009, 2017). However, especially in natural systems, covariance of traits and fitness may arise in a variety of ways, and some kinds of environmentally-induced covariance lead to estimates of *β* that poorly reflect how traits influence fitness. Selection gradients estimated in this way consequently lead to inaccurate predictions of evolution when used in the Lande equation.

Inaccurate evolutionary predictions arise because key assumptions underlying the Lande equation are often violated in empirical applications. In particular, the Lande equation makes the strong assumption that trait-fitness associations arise entirely due to causal effects of the measured set of traits on fitness. Violations of this assumption have been described in different ways: for example, as the environment “biasing” estimates of selection (Rausher 1992, Stinchcombe et al 2000), “short-circuiting” natural selection (Kruuk *et al*. 2003), or generating “danger” in for evolutionary predictions (Morrissey *et al*. 2010). These discussions centre on an important question: When and how can we reliably predict the course of adaptive evolution? However, while the above metaphors are each apt in their own way, the talk of “short-circuiting” and “danger” in particular may foster confusion about whether and how environmentally induced variation in traits and fitness can be accommodated by key approaches in evolutionary quantitative genetics, including the Lande equation. Additionally, all these discussions have been presented very much from the perspective of statistical genetics, and have so far yielded solutions to the problem of environmental confounding that are based on correspondingly complex statistical procedures. Such solutions make enormous demands on data, as they require estimating the genetic basis of the relationship between traits and fitness (reviewed in Morrissey *et al*. 2010). Despite the challenges of empirical application, the few existing attempts to test for bias due to environmentally induced covariance between traits and fitness suggest that the phenomenon might be rather common (Rausher and Simms 1989, Stinchcombe *et al*. 2002, Kruuk *et al*. 2002, Sheldon *et al*. 2003, Morrissey *et al*. 2012, Bonnet *et al*. 2017).

In this article, we first delimit the type of environmental variation that interferes with the application of the Lande equation and its close analogue, the breeders’ equation. Our analysis is rooted in evolutionary ecology, rather than the statistical quantitative genetics of previous studies. We then partition the selection differential into evolutionarily active and inactive components by (a) extending the standard decomposition based on direct and indirect selection gradients, and (b) invoking the explicitly causal concept of extended selection gradients. The latter leads to a novel and general decomposition of the selection differential, according to extended selection gradients, into total causal and incidental covariances. These decompositions elucidate the fundamental nature of the environmental covariance problem and inform the design of data collection and statistical analyses to solve the problem. Discussing environmental confounding from the perspective of phenotypic selection analysis, rather than statistical genetics, yields a simpler and more intuitive conceptualisation of the underlying problem. Importantly, our approach also calls for more straightforward – and more easily interpretable – data collection and analyses to tackle the so-called “environmental covariance problem”.

### Path diagram-based description of environmental confounding of selection measures

Figure 1 illustrates a scenario where there is environmental confounding of a typical phenotypic selection analysis. The trait (for simplicity, a single trait, though see appendix 3 for a treatment with a multivariate phenotype and multiple environmental variables) covaries with fitness for two reasons. The trait has an effect on fitness (*b*_*z→w*_), and additionally, the trait covaries with fitness because the trait and fitness share the confounding environmental variable as a common cause (i.e., because of non-zero 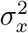, combined with non-zero *b*_*x→z*_ and *b*_*x→w*_). This scenario was considered by Scheiner *et al*. (2002), and here we dissect it in much greater detail. Rausher (1992) presented a similar path diagram in the first serious attempt to dissect the environmental covariance problem. However, a key difference between his diagram and our treatment here is that we treat the environmental confounder, *x*, as an explicit variable. Rausher (1992) treated environmental covariance as latent quantity. His treatment is entirely capable of reflecting the covariance structures represented in our figure 1, but we shall see that considering what environmental variables might be responsible for this covariance leads to a different class of solution to the problem.

**Figure 1:**
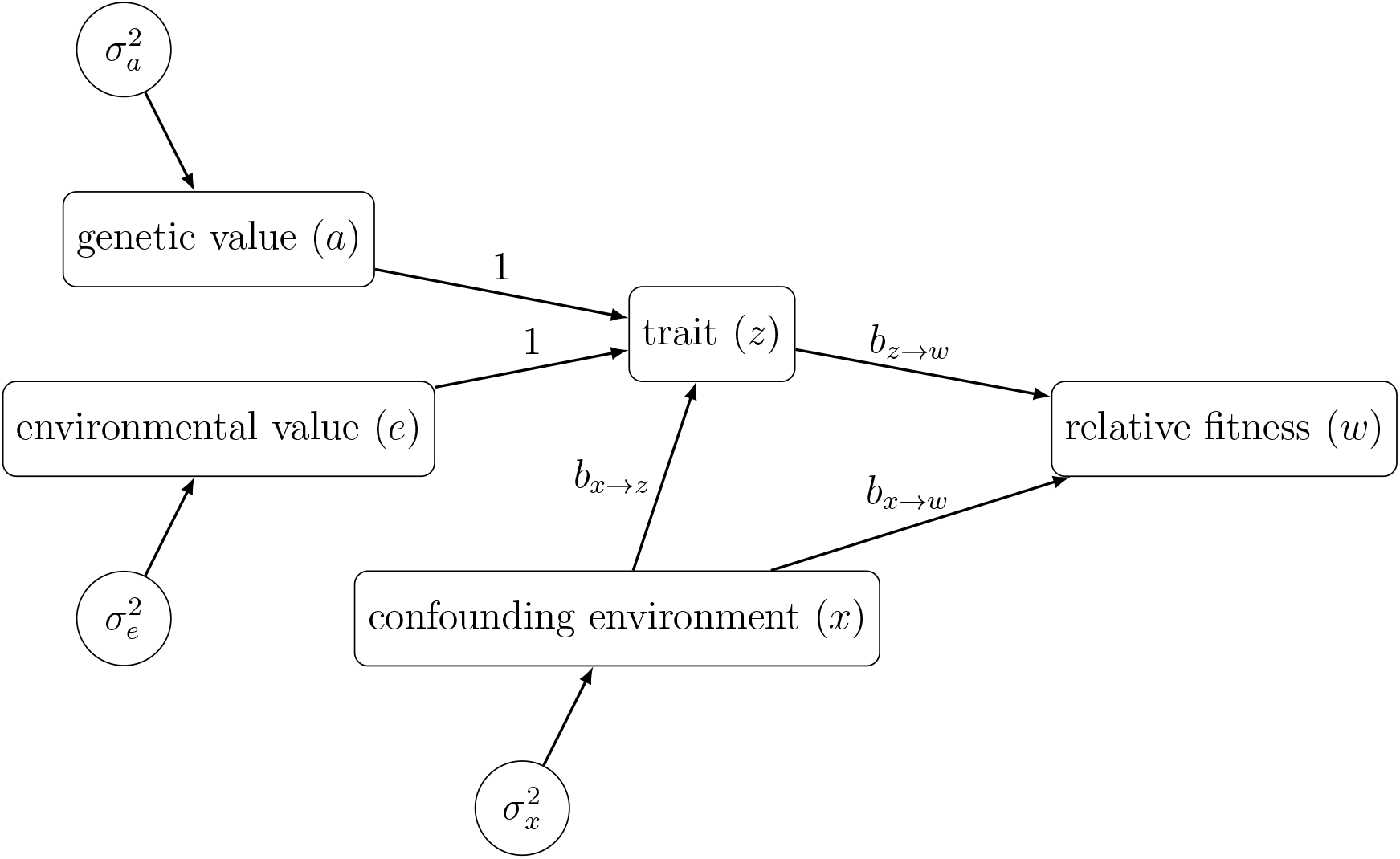
Relationships between a trait, an environmental variable, and fitness, to illustrate the essential elements of the problem of environmental confounding of estimates of selection. Terms are defined in table 1, along with key quantities that are determined by terms and relationships in the path model.

To illustrate that a standard phenotypic selection analysis goes wrong in figure 1’s scenario, we can first work out an expression for the expected evolutionary change in the trait. From the secondary theorem of selection (STS: Robertson 1966, Walsh and Lynch 2018), the evolutionary change in the trait *z* is given by the covariance of breeding values for *z* with relative fitness, i.e.,

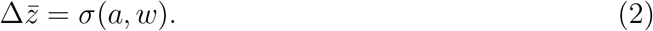

From the rules of path analysis for linear and additive effects, the covariance of any quantity *A* with another quantity *B*, arising from the ultimate (i.e., potentially mediated by intermediate variables) effect of *A* on *B* is given by the variance of *A* times the total effect of *A* on *B*. This rule, applied to the effect of the genetic value *a* on relative fitness *w*, will give the total covariance of *a* with *w*, simply because the causal effect of *a* on *w* is the only path connecting these variables. The total effect of *a* on *w* is 1 · *b*_*z→w*_, so

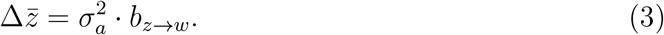

This expression should be quite familiar, as it is the Lande equation (equation 1) in a univariate form (later, we confirm that the reasoning applies in the multivariate case as well). Many readers may recognize that *b*_*z→w*_ is simply a selection gradient, except that we have placed it within a path model that allows us to elaborate on how specific measurable features of the environment influence traits and/or fitness.

To show how the scenario in figure 1 illustrates confounding of phenotypic selection analyses, consider what a classic selection analysis would estimate for the selection gradient (which has a true value of *b*_*z→w*_). Regressing relative fitness on the trait *z*, as in the classical approach, yields a selection gradient estimate of

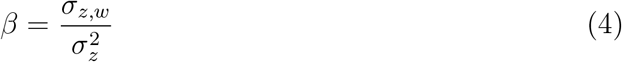

This is the ordinary least squares estimate of the simple regression coefficient, which equals the covariance of the trait and relative fitness divided by the variance of the trait. Expressions for *σ*_*z,w*_ and 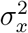 are given in table 1, and explanations of how they are obtained from the rules of path analysis are given in appendix 1. From these expressions, we can calculate the selection gradient that would be estimated for the scenario in figure 1:

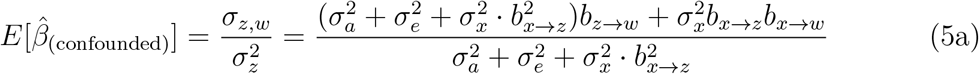

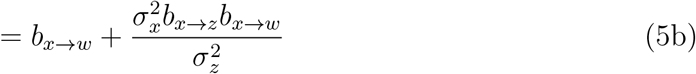

Clearly, 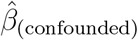 is a biased estimate of the true value of the selection gradient whenever any environmental variable exists 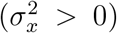 that influences both the trait and fitness (*b*_*x→z*_*b*_*x→w*_ *≠* 0).

**Table 1:** Definitions of quantities in figure 1, and formulae for some of these quantities derived using the rules of path analysis.

| quantity | symbol | formula |
| --- | --- | --- |
| trait | $z$ | |
| environmental variable | $x$ | |
| (relative) fitness | $w$ | |
| (additive) genetic variance of the trait | $\sigma_a^2$ | |
| environmental variance of the trait,<br>conditional on any explicitly modelled<br>environmental variables | $\sigma_e^2$ | |
| variance of an explicitly modelled en-<br>vironmental variable | $\sigma_x^2$ | |
| effect of the trait on fitness | $b_{z \rightarrow w}$ | |
| effect of the environmental variable on<br>the trait | $b_{x \rightarrow z}$ | |
| direct effect of the environmental vari-<br>able on fitness | $b_{x \rightarrow w}$ | |
| variance of the trait | $\sigma_z^2$ | $\sigma_a^2 + \sigma_e^2 + \sigma_x^2 b_{x \rightarrow z}^2$ |
| covariance of the trait and fitness | $\sigma_{z,w}$ | $(\sigma_a^2 + \sigma_e^2 + \sigma_x^2 b_{x \rightarrow z}^2) b_{z \rightarrow w} + \sigma_x^2 b_{x \rightarrow z} b_{x \rightarrow w}$ |
| covariance of the environmental vari-<br>able and the trait | $\sigma_{x,z}$ | $\sigma_x^2 b_{x \rightarrow z}$ |
| covariance of the environmental vari-<br>able and fitness | $\sigma_{x,w}$ | $\sigma_x^2 (b_{x \rightarrow w} + b_{x \rightarrow z} b_{z \rightarrow w})$ |

The environmental confounding of selection illustrated in figure 1 is the same essential phenomenon discussed in prior literature (Rausher 1992, Kruuk *et al*. 2003, Morrissey*et al*. 2010). However, we hope that putting it into the more tangible framework of a path diagram, rather than the involved machinery of statistical genetics, provides a more accessible description of the problem.

### Multiple regression estimates of *β* that account for confounding (environmental) variables

Figure 1 motivates a very simple analysis to estimate the direct effects of traits on fitness (i.e., selection gradients) in the presence of confounding by environmental variables. Just as multiple regression can be used to estimate the direct effects of each of a set of traits on fitness (i.e., the classic Lande-Arnold analysis), it can also isolate such direct effects while accounting for potentially confounding environmental variables. The key is to simply include both individual trait values *and* individual-level measures of environmental variables as predictors in the multiple regression. Biologists use multiple regression analysis for exactly this purpose all the time. Strangely, however, this elementary approach is not routinely used in studies of selection.

Here, we verify that multiple regression leads to correct estimates of selection gradients when fitness is regressed simultaneously on trait values and environmental variables. The partial regression slopes from the multiple regression of relative fitness on a trait *z* and an environmental variable 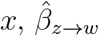 and 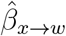 respectively, are given by

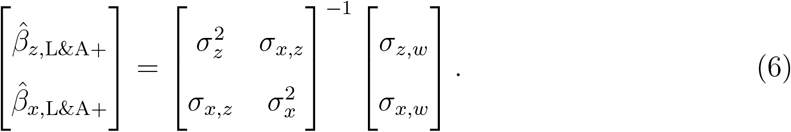

The estimates 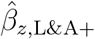 and 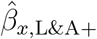 correspond to selection gradients in the classical approach of Lande and Arnold (1983), except that both the trait and the environmental variable are treated as predictors in the regression (i.e., as ‘traits’ in the classical terminology). Note that equation 6 is the multivariate analog of the expression for the regression slope in the equation 4 (i.e., it is the inverse of the covariance matrix of predictors multipled by the vector of covariances of the predictor variables with relative fitness). Isolating the selection gradient estimate we are most interested in, i.e., 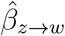, we can expand this equation to obtain

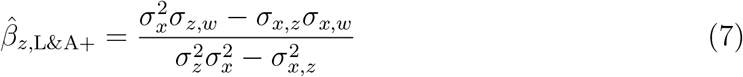

Substituting the expressions for component terms given in table 1, and simplifying, yields 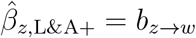 (see appendix 2 for intermediate steps). For some readers, this result will be intuitively obvious: The path coefficient describing the effect of *z* on *w* is estimable as the partial regression coefficient of *w* on *z*, controlling for *x*. However, the formal clarification in appendix 2 might be useful to anyone less aware of the close relationship between multiple regression and path analysis. Similarly, there seems to be substantial, but misplaced, concern among biologists about the ability of multiple regression to discern direct effects among correlated predictors (reviewed in Morrissey and Ruxton 2018), and the present situation is inherently one involving correlated, or collinear, predictors. Thus, the confirmation that 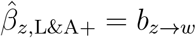 in appendix 2 may also help to alleviate any concerns as to whether the correlation structure inherent to the environmental confounding problem is well-handled by multiple regression analyses involving both traits and environmental variables as predictors. Appendix 4 gives example code for numerical demonstrations that correct evolutionary prediction (i.e., agreement with the STS) can be made using estimates of selection gradients obtained from extensions of the Lande-Arnold approach to include environmental variables.

### Partitioning the selection differential in the presence of confounding (environmental) variables

So far, we have considered quantitative measurement of selection, and the potential for such measures to be confounded by environmental variables, via the selection gradient, *β*. However, the basic idea applies equally to the selection differential, *S*, which is the covariance of a trait with relative fitness. The selection differential is grounded in evolutionary theory via an analogue of the Lande equation, the breeder’s equation (Falconer 1981)

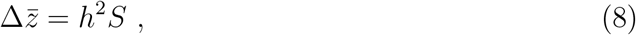

where *h*^2^ is the heritability, or the proportion of the phenotypic variance of a trait that arises due to additive genetic differences among individuals. Just as we saw that a standard estimate of *β* (from regression of *w* on *z* only) does not correctly predict evolution – recall that equation 5 did not produce the selection gradient estimate needed for equation 3 to work correctly – a standard calculation of *S* does not correctly predict evolution using the breeders’ equation. Table 1 gives the covariance of trait and fitness that would provide an empirical estimate of *S*, which, used in the breeders’ equation, would predict an evolutionary change due to selection of

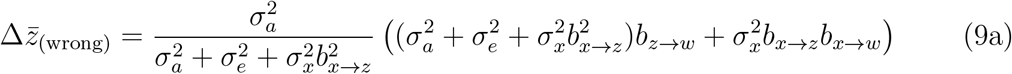

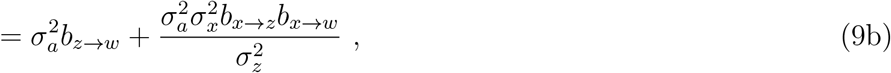

which does not agree with the STS and is therefore wrong. Note that we have simply reproduced the confounded estimate of *β* (equation 5), only scaled by 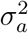. Thus, whether viewed through selection gradients or differentials, environmental confounding has the same ultimate effect on predicting evolution. The rearrangement in part (b) expresses the erroneous prediction of the breeders’ equation in two terms, the first of which is a correct version of the Lande equation (it coincides with equation 3). Therefore, the breeder’s equation is correct when the second term is zero due to any of the conditions that remove the confounding path from figure 1, namely if 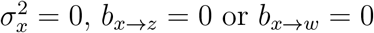.

We now further dissect *S* under the conditions of figure 1 to gain more precise insight into how confounding derails predictions made using the breeders’ equation. We thereby clarify which aspects of environmental variation are accommodated by standard analyses, and which cause problems. In turn, this leads to intuitive ways to make use of ‘environmentally informed’ regression-based estimates of natural selection based on *β*_L&A+_. Based on the path diagram (figure 1), we can decompose *S*, the covariance of the trait and fitness, into four components. These are:

a. covariance between the trait and fitness that ultimately arises from genetic variation for the trait 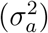,
b. covariance between the trait and fitness that ultimately arises from environmental effects on the trait 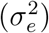, other than those generated by the explicitly-treated environmental confounder *x*,
c. covariance between the trait and fitness that arises from effects of the environmental variable *x* on the trait, combined with the fact that the trait affects fitness, and
d. covariance arising from the dual effects of the environmental variable on the trait and on fitness.

The first three components relate to the causal path *z* → *w*, with the total covariance via this path split up according to the various sources of variance in *z*. The last component corresponds to the confounding path *z* ← *x* → *w*. This decomposition consequently accounts separately for all exogenous sources of variance, and all ways that such sources contribute to covariance of the trait with fitness. Corresponding to this decomposition, we can re-write the expression for *σ*_*z,w*_ in table 1 as follows:

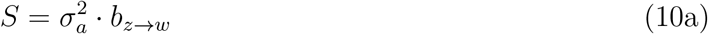

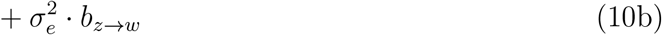

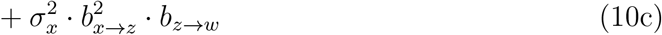

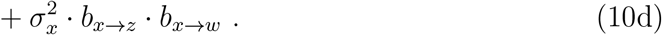

As above, we can again see that if any of 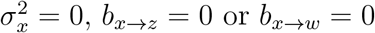 or *b*_*x→w*_ = 0, then part (d) of equation 10 makes no contribution to *S*, and the resulting value of *S*, when entered into the breeder’s equation, generates an evolutionary prediction that coincides with the STS (and is therefore correct):

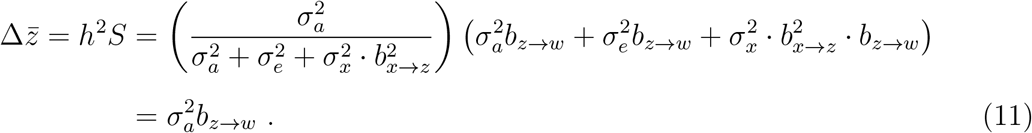

Recall that this last expression is the formula obtained by applying the STS to figure 1 (equation 3).

That only the environmental contribution in equation 10d needs to be eliminated for the breeder’s equation to accurately predict evolutionary change is important to understand. Equations 10b and c also represent contributions of environmental variation to *S*, but these environmental contributions to *S* do not invalidate the breeder’s or Lande equations. This accommodation of environmental variation applies both to environmental effects that are not explicitly treated, but rather treated as latent variability 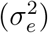, and to variation that is explicitly incorporated into a conceptual model (i.e., variation in the explicit environmental variable *x* in figure 1). It is only that component of environmental covariance between a trait and fitness that does not involve the causal effects of the trait on fitness that generates estimates of selection differentials and gradients that give wrong evolutionary predictions.

We typically think heuristically that a non-zero selection differential *S* and positive heritability *h* will lead to evolutionary change. Importantly, however, any part of *S* that is not generated by direct or indirect selection on traits is evolutionarily inert. Regardless of what the breeder’s equation may predict, environmentally induced covariance between a trait and fitness that does not arise from the causal effects of traits on fitness does not generate an evolutionary response. The metaphor of Kruuk *et al*. (2003) is thus quite fitting: selection may be partly null – or “short-circuited” – in that it will not generate the evolutionary change we would expect based on the breeder’s equation. A more mechanistic characterisation of the situation might be to recognise that the selection differential represents covariance between a trait and fitness arising via a variety of processes. Only some of these processes generate evolutionary change, even though all are typically referred to as “selection”.

Given coefficients of the regression of relative fitness on traits and environmental variables, we can factor the *b*_*z→w*_ term in equations 10a-c in a way that decomposes the selection differential into two parts, rather than four. The first part represents the component of *S* that works in the breeder’s equation according to the reasoning given in equation 11. The second part is the component that confounds inference of selection, and thus generates erroneous predictions of evolution. The two-part decomposition is given by:

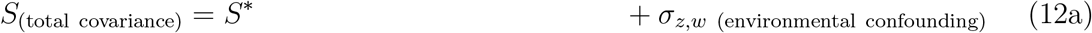

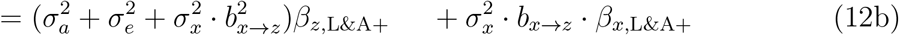

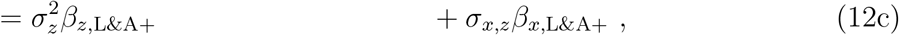

noting that 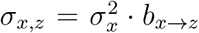 (table 1). *S*^*∗*^ is that component of the selection differential that would give the correct evolutionary prediction when substituted into the breeder’s equation, given a good estimate of the heritability, under the standard assumption that there is no direct selection on other correlated traits (Walsh and Lynch 2018). This is mathematically analogous to the standard decomposition of the selection differential into direct selection (equal to the selection gradient of the focal trait multiplied by the variance in that trait) and indirect selection (equal to the sum of selection gradients of all other traits, weighted by their covariance with the focal trait). This construct is merely extended to define analogues of selection gradients for environment-fitness effects (*β*_*x*,L&A+_) and to separate these effects from those of phenotypic traits.

Up to this point, we have focused on analyses with a single phenotypic trait. This was mostly to avoid scary matrices (but these *are* coming). However, as the relationships between selection differentials and gradients are most meaningful in multivariate selection analyses, it is worth giving the full decomposition of *S* in the case of *m* traits, *z*_1_, …, *z*_*m*_ and *n* environmental variables *x*_1_, …, *x*_*n*_. The multivariate equation, without matrices, but for only a focal trait *z*_1_, is

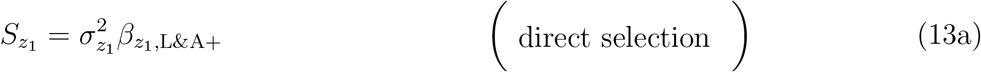

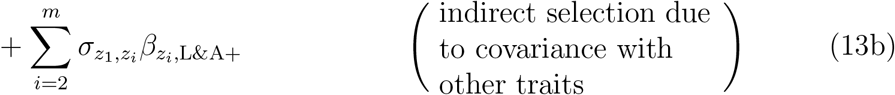

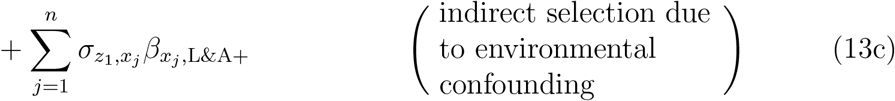

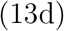

In a more compact and complete presentation, the vector of selection differentials for traits of interest depends of the covariance matrix of the traits **P**, the vector of selection gradients for those traits ***β***_L&A+_, the covariances of the traits with the environmental variables **Σ**_***x***,***z***_, and the vector of environmental analogues of selection gradients ***β***_*x*,L&A+_

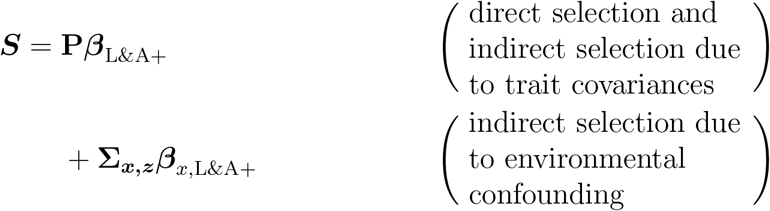

Without going to lengths in unpacking these multivariate decompositions of the selection differential vector, there is really just one key elaboration in the multivariate case that is not immediately apparent in the univariate treatment: Even if no single environmental variable influences both a trait and fitness, environmental confounding can still occur if, among a correlated set of environmental variables, some influence traits and some influence fitness. This is encapsulated in the fact that the environment-fitness effects 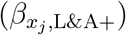 in equation 13c contribute to *S* via trait-environment covariances *σ*_*z,x*_. These trait-environment covariances may arise from any causal structure, including direct effects of a specific environmental variable on a focal trait and indirect associations among environmental variables.

### More general path models

The direct effects of environmental variables on fitness, combined with their effects on traits, are responsible for the environmental covariance problem. In the absence of direct effects of the environment on fitness, which can be estimated and corrected for using multiple regression, environmental covariance does not confound phenotypic selection analysis and evolutionary predictions. Therefore, while path analysis was useful to clarify the environmental covariance problem and justify the multiple regression-based solution, path analysis is not strictly necessary for empirical studies to use environmental data to directly address the environmental covariance problem. However, considering the problem in the context of more general path analysis-based phenotypic selection analysis has two benefits. First, we can address the decomposition of the selection differential using the extended selection gradient concept (Morrissey 2014, 2015, Henshaw *et al*. 2020), which will often provide more biologically informative summaries of how traits influence fitness (Scheiner *et al*. 2000). Extended selection gradients potentially fully reflect how traits influence fitness, accounting for both direct and mediated effects, and as such can reflect Sober’s (1984) distinction between selection ‘of’ and selection ‘for’ a trait. Second, this more general analysis yields additional insights into the environmental confounding problem that are not apparent from the regression analyses presented so far.

Figure 2 represents the key elements of the general case:

**Figure 2:**
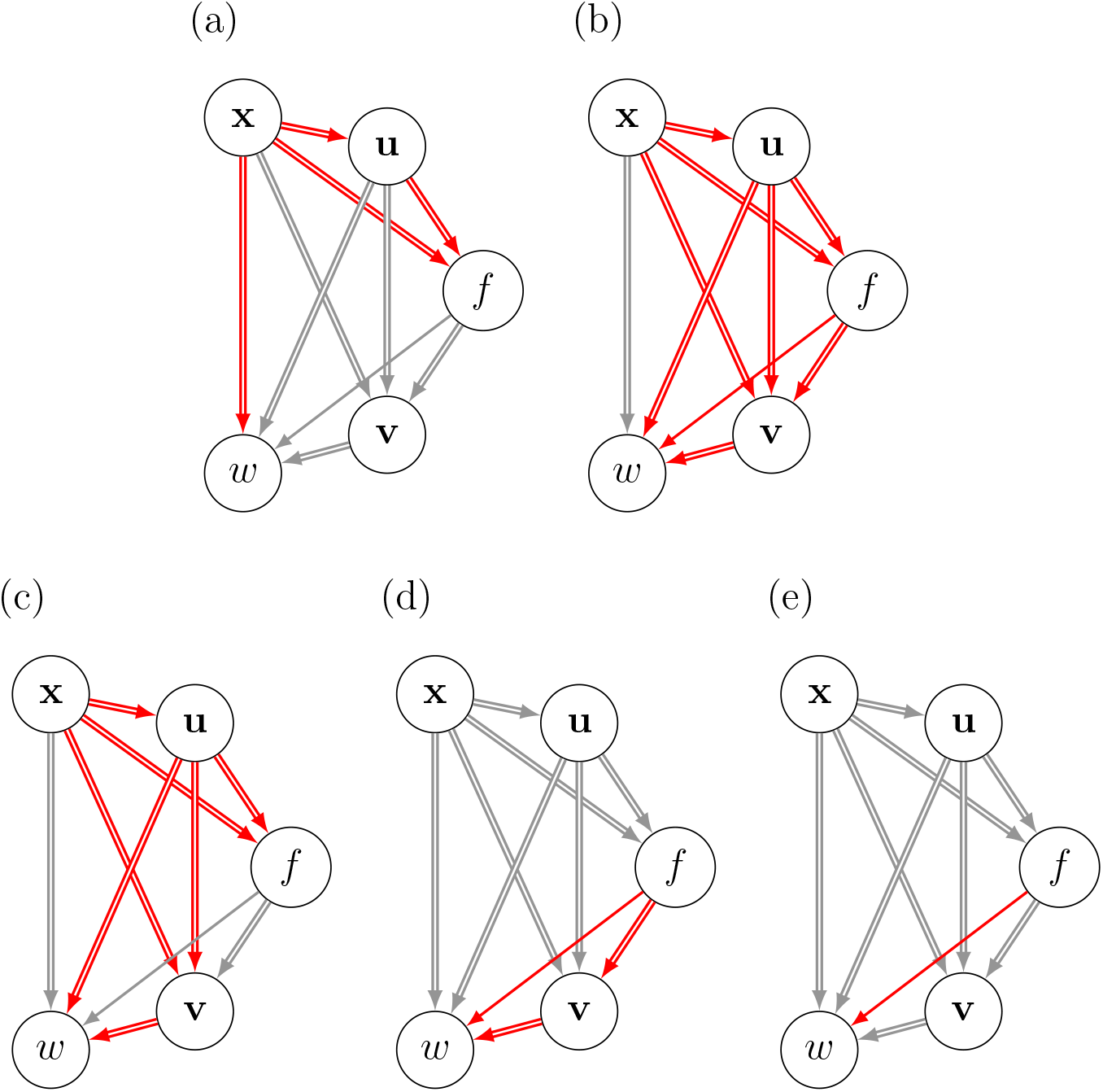
Path diagrams relevant to the decomposition of the selection differential *S*_*<sub>f</sub>*_ of a focal trait *f*, considering potential environmental variables (**x**), traits causally preceding the focal trait (**u**), and traits mediating ultimate causal effects of the focal trait on fitness (**v**). Double-lined arrows potentially represent multiple causal effects among one or more of the vector-valued nodes. Red arrows represents sets of paths that contribute in conceptually distinct ways to covariance of the focal trait with fitness: (a) paths that can generate environmental confounding of selection on the focal trait, i.e., environmentally induced covariance that causes erroneous evolutionary predictions if not accounted for; (b) paths potentially contributing to *S*_*<sub>f</sub>*_ that do not confound applications of the breeders’ or Lande equations, even if the environmental variables **x** are not considered explicitly; (c) paths generating incidental covariance of the focal trait with fitness (limited to those paths allowing correct evolutionary prediction without explicitly considering **x**); (d) paths that can generate causal association of the focal trait with fitness, as quantified via the extended selection gradient *η*_*<sub>f</sub>*_ ; and (e) the direct effect of the focal trait on fitness, as quantified by the direct selection gradient, *β*_*<sub>f</sub>*_ (n.b., only this path is represented by *β*_*<sub>f</sub>*_, regardless of whether *β*_*<sub>f</sub>*_ is estimated by path analysis or multiple regression).

- environmental confounding variables, **x**
- traits, **u**, that causally precede a focal trait, *f*
- more traits, **v**, that may depend on **x, u** and *f*, and
- fitness, *w*.

Arrows with double lines represent the potential for multiple effects, e.g., the double arrow from **x** to *f* indicates that multiple environmental variables might influence the focal trait. Similarly, the double arrow from *f* to **v** indicates that the focal trait, *f*, may influence multiple other traits. Each sub-figure in figure 2 illustrates the analogue of a component of covariance of the focal trait with fitness, but expanded to the multivariate case with explicit effects of traits on one another. Critically, we will now be working with extended selection gradients (Morrissey 2014, 2015, Henshaw *et al*. 2020), which represent the total effect of traits on fitness (e.g., those effects corresponding to the paths *f* → *w* and *f* → **v** → *w* in figure 2d), rather than direct selection gradients (e.g., the effect arising via the single path *f* → *w* in figure 2e).

The covariances among environmental variables **x** and traits **z** = [**u**^*T*^, *f*, **v**^*T*^]^*T*^ specified by the path model in figure 2 may be written

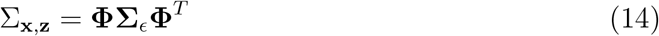

where **Φ** contains the total (rather than direct, as in the *b* coefficients in figure 1) effects of traits on one another (see Morrissey 2014 for calculation of **Φ** from a corresponding matrix of *b* coefficients). We can write **Φ** in block form to highlight its structure:

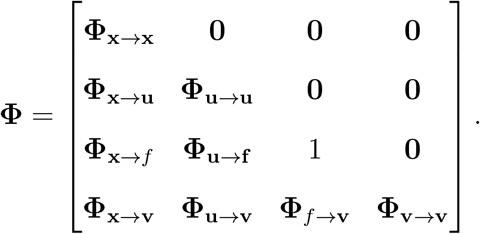

Each single or double arrow in figure 2 is represented by a corresponding element of the **Φ** matrix, representing the total causal effects of any one element of the path model on any other. Additionally, for generality, we accommodate the fact that causal relations may exist among elements within each of the vector-valued quantities, **x, u**, and **v** (e.g., one environmental variable may influence another). The corresponding terms are given by the matrices along the (block) diagonal of **Φ**. Note that the focal trait’s influence on itself is simply 1. Variation in and associations among environmental variables, traits, and fitness ultimately arises from exogenous inputs of variance to the system, which are

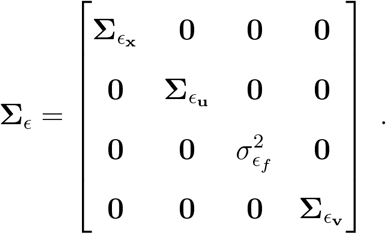

The vector of selection differentials (i.e., covariances of traits with relative fitness) is given by the matrix product of the covariance matrix of the traits **x** and **z** with the vector of selection gradients ***β***_**x**,**z**_, namely **S** = Σ_**x**,**z**_***β***_**x**,**z**_, which yields expressions for all covariances of elements of **x** and **z** with relative fitness. Focusing on the focal trait *f*,

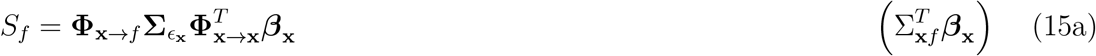

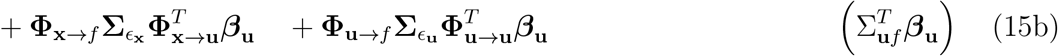

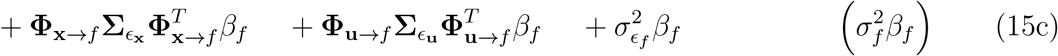

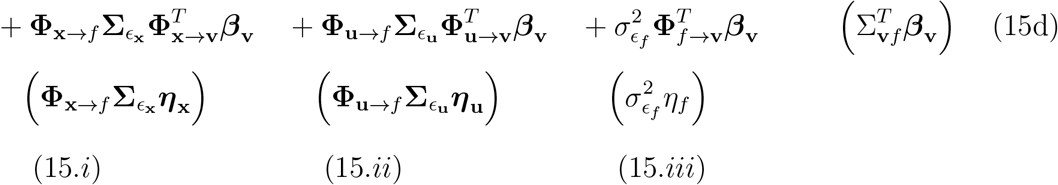

Quantities in parentheses down the right side and across the bottom are not part of the equation. Rather, the terms in each row of the equation (labelled 15a-d) sum to the parenthetical expressions on the right-hand side, which represent a decomposition of the selection differential in terms of the direct selection gradients (i.e., the decomposition that would be given by Lande and Arnold (1983)). The equation for *S*_*f*_ breaks down each component of the Lande-Arnold decomposition further according to the different sources of covariance between the focal trait and correlated traits and environments. Similarly, the terms in each column (labelled 15*i* -*iii*) sum to the parenthetical expressions at the bottom of equation 15. These collumn-sums represent a decomposition of the selection differential according to extended-sense selection gradients, or total effects of traits on fitness (Morrissey 2014, 2015, Henshaw *et al*. 2020).

The partition of the selection differential according to extended selection gradients has not previously been treated, and it has some interesting features that complement the well-known partition based on direct selection gradients. Whereas direct selection gradients link the total variances and covariances in and among traits to the selection differential, extended selection gradients, such as *η*_*f*_, link independent inputs of variation to focal traits (e.g., 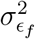) to the selection differential. Loosely, extended selection gradients partition *S* according to the ultimate sources of variation in a system where traits affect one another and fitness. Importantly, the total fitness effect of a trait includes indirect effects passing through other traits (see more extensive descriptions in Morrissey 2014). Traits that are causally ‘downstream’ of a focal trait (i.e., **v** in figure 2) consequently do not figure explicitly in the above decomposition of the selection differential, but are rather included in the extended selection gradient *η*_*f*_ on the focal trait. This is why, given the causally ordered sets of traits, **x, u**, *f* and **v**, there are four elements in the decomposition according to direct selection gradients (eqs. 15a-d), but only three elements in the decomposition according to extended selection gradients (eqs. 15*i* -*iii*).

The decomposition of *S*_*f*_ based on the extended selection gradients is expressed in terms of the exogenous variance of each variable. Exogenous inputs are latent quantities – variation over and above that attributable to the modelled causes of a variable – and so it might be more intuitive to re-formulate the decomposition in terms of the total tangible variance of a focal trait. It is possible to express the partition of *S*_*f*_ partially in terms of the variance of a focal trait, yielding a decomposition that includes a simple form for the contribution of a trait itself to its own selection differential, which is 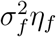. This total causal covariance has a component represented by the familiar contribution of direct selection, 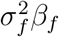. The other component, 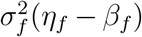 or 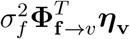, represents causal effects of *f* on fitness that are mediated by other traits, **v**. This decomposition into causal and incidental components of *S*_*f*_ is shown in equations A2 to A7 in appendix 3, which re-arrange equation 15 as follows:

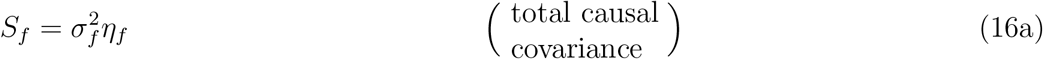

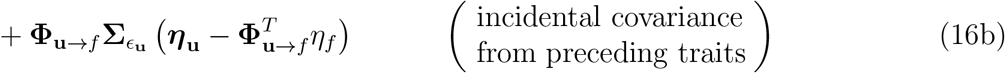

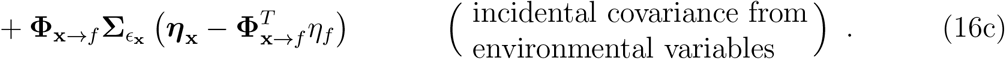

The total incidental association is just the difference between *S*_*f*_ and its total causal component 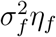. However, if the incidental covariance is to be decomposed into contributions from individual traits (or groups of traits and environments as is the case in equation 16) the expressions are more complicated. Partitions of the incidental covariance cannot be expressed as simple functions of covariances among traits, as are indirect effects in the decomposition using direct selection gradients. Instead, the decomposition of incidental effects using extended selection gradients is governed by the association of covariance inputs for **x** and **u** with 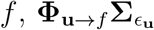 and 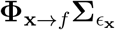, respectively, and the the parts of the total effects of **x** and **u** on fitness that are not mediated by *f*, which are 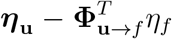 and 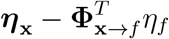, respectively.

The decompositions of the selection differential in equations 15 and 16 requires the linearity and additivity of all effects (i.e., the effects of environmental variables on traits, of traits on one another, and of environmental variables and traits on fitness). More generally, extended selection gradients can be defined and used as the basis for multivariate evolutionary predictions even in the presence of arbitrary non-linear and non-additive effects (Morrissey 2015, Henshaw et al. 2020). In appendix 5, we accordingly present a more general form of the above decomposition that allows for non-linear and non-additive effects. This more general form assumes only the normality of the focal trait. It is not clear that an additive partition of *S* is possible for a non-normal focal trait given arbitrary non-linear causal effects among traits.

Consideration of the full multivariate situation raises two considerations for the use of measured environmental variables in phenotypic selection analyses. First, as we noted in our multivariate treatment based on direct selection gradients, if environmental variables are correlated, either causally or for reasons that are not explicitly modelled, then environmental confounding can occur even if no single environmental variable influences both a trait and fitness. This is illustrated in figure 3a, where the environment induces an association between the trait and fitness, even though no single environmental variable influences both quantities. In this situation, the confounding can be eliminated by measuring either environmental variable. This suggests that judicious consideration of environmental variables can sometimes lead to uncoufounded inference of selection even if it is not possible to measure all environmental variables that are causally relevant to trait-fitness associations. Second, it is possible to use traits to address environmental confounding, without explicitly measuring environmental variables, in some circumstances. Consider figure 3b. Since all covariance of *z*_2_ and *w* contributed by *x* acts via *z*_1_, a standard multivariate analysis of direct or extended selection gradients will be confounded for *z*_1_, but not for *z*_2_. Absent consideration of *x*, the direct selection gradient vector will be partly confounded, and thus will not generate correct evolutionary predictions using either the Lande equation or its analogue based on extended gradients given in Morrissey (2014). However, this does not mean that all elements of ***β*** and ***η*** are confounded. In other words, correct ecological interpretations of some selection gradients may be possible in the presence of environmental confounding, provided that the confounding is mediated by preceding traits (which themselves will not have estimated selection gradients properly represent their effects on fitness). Pearl (2009) provides a complete treatment of which sets of variables can be controlled for in order to correctly estimate causal effects in the presence of confounding (see also Henshaw *et al*. 2020).

**Figure 3:**
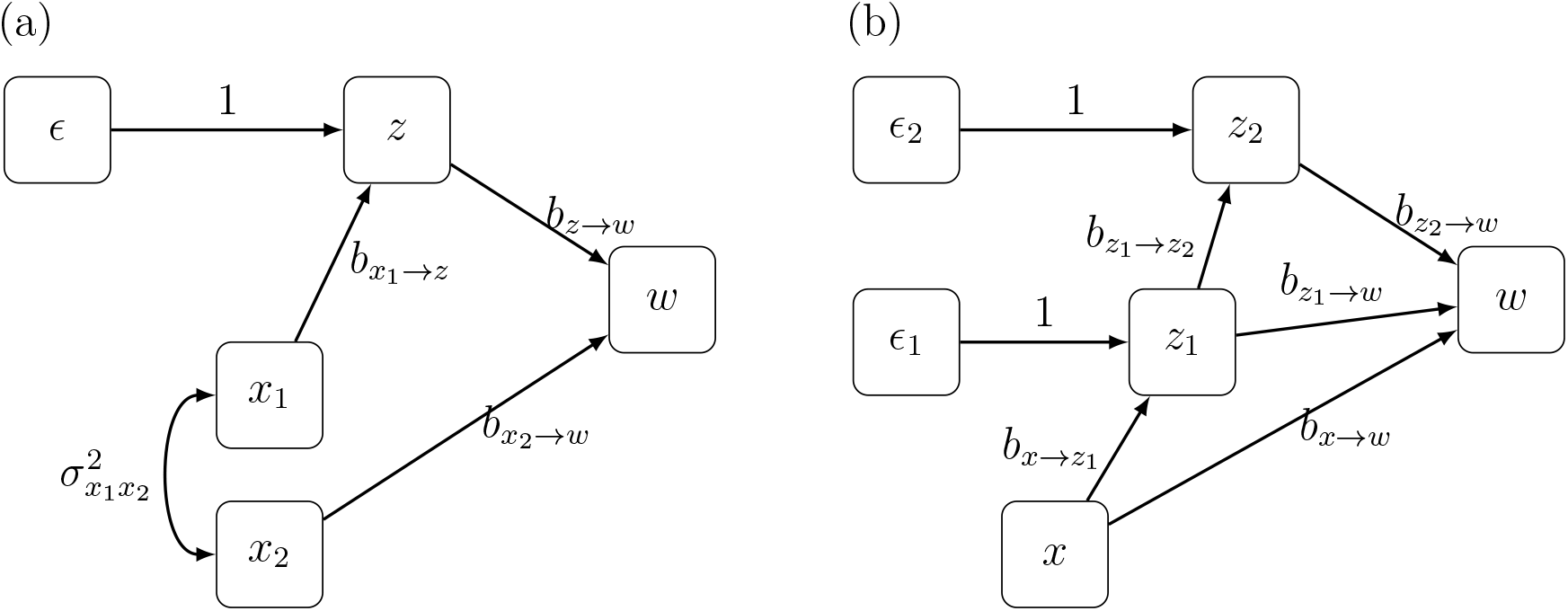
Cases where it is not necessary to measure and model all variables that generate trait-fitness covariance in order to generate unconfounded estimates of selection gradients. In (a), inclusion of either environmental variables *x*_1_ or *x*_2_ in an analysis of direct or extended selection gradients will generate correct estimates of selection for the trait, *z*. In (b), inclusion of *z*_1_ allows correct estimation of selection for *z*_2_, even though the environmental variable *x* ultimately causes spurious covariance of *z*_2_ with fitness. Note that in both cases, the variables mentioned constitute ‘backdoor sets’, controlling for which allows one to isolate the causal effect of the focal trait on fitness (for details see Pearl 2009; Henshaw *et al*. 2020).

## Discussion

We hope that these discussions of environmental confounding will clarify the nature of the problem and its implications for selection analysis. Rather than a potentially nebulous aspect of environmental trait-fitness association that must be tackled with complex quantitative genetic analyses, we have shown that the phenomenon can be understood straightforwardly using multiple regression – the standard method for disentangling direct and indirect effects. From a practical perspective, our suggestion represents a modest extension of the framework of Lande and Arnold (1983), which is already familiar to many evolutionary biologists. This solution has the advantage of retaining the standard estimation procedures for key evolutionary parameters, while potentially improving the quality and biological interpretability of such estimates. This more standard solution thus potentially yields more biological insight than the statistical genetics solutions that have been presented to date as the main solutions to the ‘environmental covariance problem’.

Empirical biologists are frequent users of multiple regression, using it in a wide range of contexts that go far beyond estimating selection gradients via Lande and Arnold’s (1983) method. Why, then – despite decades of discussion about environmental confounding of selection gradient estimates (Rausher 1992, Kruuk *et al*. 2003, Morrissey *et al*. 2010) – has multiple regression-based selection analysis hardly been used or even discussed as a solution? First, the problem may be that environmental confounding has typically been treated from the perspective of very involved statistical genetics approaches. This may have obscured the simple nature of the problem. To this end, perhaps simply depicting the problem graphically as a standard instance of a confounder variable (figure 1), and then dissecting in detail what that means for the application of familiar multiple regression analyses, will already help to clarify which standard statistical approaches might potentially be employed to combat the problem.

Another potential impediment to incorporating environmental variables into multiple regression-based selection analyses may simply be fear that any arbitrary change to the standard approach may not be permissible. Even in the absence of environmental confounding, selection analyses must be constructed in particular ways in order to generate correct values of *β* (in the sense that they will correctly predict evolutionary change). The requirements for constructing regression-based selection analysis thus go well beyond the normal requirements of multiple regression itself as a general statistical procedure. For example, the response variable must be relative fitness (Lande and Arnold 1983), and fitness proxies may not generally be used (Franklin and Morrissey 2017). Similarly, predictor variables need to be entered in specific ways, especially with quadratic terms scaled appropriately (Stinchcombe *et al*. 2008). As such, any given arbitrary elaboration of regression models of selection – while perhaps consistent with the requirements of multiple regression analysis in general – may not necessarily still return valid selection gradient estimates. Consequently, it may not previously have been clear that it is ‘permissible’ (i.e., producing correct selection gradients that quantitatively predict evolution) to include environmental variables in regression-based analyses of natural selection.

Finally, because the key theory of selection analysis (e.g., Lande and Arnold 1983, Arnold and Wade 1984, Wade and Kalisz 1989) is presented in a way that does not specifically consider environmental confounding, all of the key theoretical articles that have motivated empirical selection analyses have only written about trait-fitness relationships. Consequently it is not surprising that researchers have not been specifically motivated to characterise environmental variables that are likely to confound estimates of trait-fitness relationships, especially given the already large investment of effort to measure traits and fitness in order to conduct standard studies of natural selection.

The idea that that environmental variables could be included in the type of multiple regression analysis inspired by Lande and Arnold (1983) has arisen before in the methodological literature on selection analysis (Mitchel-Olds and Shaw 1987, Scheiner *et al*. 2002). Unfortunately, neither of these previous works have motivated researchers to measure and model potential environmental causes of trait-fitness confounding. Scheiner *et al*. (2002) considered path models similar to figure 1, but provided little justification for their (correct) assertion that including an environmental variable in a path analysis generates estimates of *β* that have the same theoretical justification. The empirical components of Scheiner et al. (2002) do not actually conduct multiple regression or path analyses that include environmental variables, but rather compare genetic and phenotypic relations among traits and fitness. We hope that our justification for including environmental variables in selection analyses will stimulate greater use of this key idea.

Mitchell-Olds and Shaw (1987) similar implied that incorporating environmental variables into regression-based analyses of selection is permissible on theoretical grounds. However, they quickly dismissed the idea that this approach could be useful in practice, because there is no guarantee that available measures of environmental variables will be suitable to entirely eliminate non-selective covariance of traits and fitness. This is of course true, but the concern is relevant to any use of multiple regression, including standard phenotypic selection gradient analysis. Any standard analysis of selection gradients will only generate estimates of *β* that reflect the direct effects of traits on fitness to the extent that a sufficiently complete set of relevant traits is included in the analysis. That a perfect set of traits may not be available does not seem to have hindered the accumulation of thousands of estimates of selection gradients.

Looking forward, our extension of the Lande-Arnold approach to explicitly include environmental variables might help motivate the design of future studies. To the extent that studies of selection are judged by their choice of phenotypic traits, the reasoning in this article suggests that equal consideration should be given to whether key environmental variables have been included to reduce the potential for environmental confounding. We consequently suggest that it could be worthwhile to invest as much effort into accounting for potentially confounding environmental variables as is typically expended in measuring multivariate phenotype.

Previous solutions to the environmental confounding problem have involved assessing the genetic basis of the the relationship between traits in fitness. Each variant of this approach is in some form an attempt to operationalise the secondary theorem of selection (Rausher 1992, Kruuk *et al*. 2003; Morrissey *et al*. 2010). A benefit of these approaches is that they do not require that confounding variables in the environment are identified, meaningfully measured, and adequately modelled. As such these more involved quantitative genetic approaches can generally be hoped to yield unbiased predictions of evolution. However, those analyses will give little insight into the reasons why traits and fitness covary, which is a primary benefit of the phenotypic selection gradient approach.

A more practical drawback of directly estimating the genetic association of traits with fitness is the extreme demands that this endeavour puts on data. In all of its forms, this approach requires immense amounts of trait and fitness data on individuals with known pedigree relationships. Except in very few study systems, limited data will mean that operationalisation of the STS will generate very noisy estimates. While it is bad to be misled by biased results, it isn’t clear that extreme uncertainty due to power-hungry analyses is much better! Pragmatically, it seems that a diversity of approaches is best for tackling the general task of understanding why traits covary with fitness, and what the evolutionary consequences of those associations will be.

## Acknowledgements

MBM is supported by a University Research Fellowship from the Royal Society (London). JMH is funded by the German Federal Ministry of Education and Research (BMBF)

## Appendix 1: Derivation of variances and covariances in the path model

Here we attempt to distill the principles of path analysis into a minimal set of rules. A more detailed treatment aimed at biologists is given in Shipley (2000). A contrast between our treatment here and most expositions (but see for e.g. Loehlin 2004, ch. 1) of these rules is that we treat the general case where variables may have arbitrary variance. Most treatments of these rules – and indeed potentially even definitions of path analysis – are presented for correlations and partial correlations, or in other words, for the case where every variable has been pre-standardised to unit variance. This approach somewhat simplifies the analysis of path diagrams, and was especially useful for simplifying expressions before computers were available (e.g., see various insightful cases in Wright (1934). While qualitative aspects of the present article should be easily accessible to users of traditional (i.e., using correlations and partial correlations) path analysis, such readers may find this appendix helpful to extend path analytical thinking to the quantitative relations presented herein.

Quantitative analysis of general (i.e., unstandardised, or in any arbitrary standardisation) path diagrams requires two sets of rules. The first set of rules governs the calculation of effects of variables on one another. The second set of rules governs the variances and covariances among traits that arise from direct or total effects.

The rules governing the calculation of effects can be summarised as follows, with the first defining a direct effect, and the second and third detailing how how direct effects are combined to obtain total effects:

1. direct effects: A direct effect is represented on a path diagram by an arrow between two variables. In figure 1, *b*_*x→z*_, *b*_*z→w*_ and *b*_*x→w*_ are direct effects, as are the effects of *a* and *e* on *z*. When a given focal variable is affected by more than variable, and those causal variables may be correlated (for example, *w* which is affected by *x* and *z* in figure 1) the direct effects are conceptually the slopes of the partial regressions, i.e., as obtained from multiple regression analysis by ordinary least squares, of the outcome variable (e.g., *w*) on its causes (*x* and *z*). In practice, direct effects can be estimated as partial regression slopes using the ‘piecewise’ approach (i.e., using a separate regression model for each outcome that is subject to causal effects) is used; similarly, alternative path model fitting algorithms aim to estimate quantities with the meaning of partial effects, but that may differ quantitatively, depending on finite data and model specification. If only one causal variable affects some outcome, or the various causes of a given outcome are known, or in practice, assumed, to be uncorrelated, then direct effects coincide with the simple (one predictor) regressions of the outcome on its cause or causes.
2. mediated effects: Effects acting along sequential paths combine multiplicatively. For example, in figure 1, the part of the total effect of *x* on *w* that is mediated by *z* is given by the product of the effects of *x* on *z* and of *z* on *w*, i.e., *b*_*x→z*_*b*_*z→w*_.
3. concurrent effects: Effects along parallel paths combine additively. For example, in figure 1 the total causal effect of *x* on *w* is comprised of the path that is mediated by *z*, and the direct path from *x* to *w*. So, the total causal effect of *x* on *w* is *b*_*x→z*_*b*_*z→w*_ + *b*_*x→w*_.

The second set of rules governs how effects, be they total, or some component of a total effect, contribute to variances and covariances among variables. These represent the basic and general mechanics of variances and covariances of linear transformations of random variables, and Wright’s general method can be thought of as the intersection of these rules with the principles above for relating direct effects to total effects:

i. if *X* has a linear effect on *Y* with slope *b*_*X→Y*_, then the contribution of *X* to variance in *Y* is 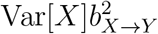
ii. if *X* has a linear effect on *Y* with slope *b*_*X→Y*_, then the contribution of this causal effect to the covariance of *X* and *Y* is Var[*X*]*b*_*X→Y*_
iii. if *Y* and *Z* are both causally dependent on *X*, with slopes *b*_*X→Y*_ and *b*_*X→Z*_, respectively, then the common cause *X* contributes Var[*X*]*b*_*X→Y*_ *b*_*X→Z*_ to the covariance of *Y* and *Z*

Elaboration of the application of these three rules to figure 1 is as follows:

- 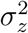 : The trait variance results in part from the sum of the latent genetic and environmental effects, with variances 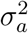 and 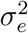, respectively. Each of these contributes directly to *z* with a slope of 1. Additionally, trait variance is contributed from the explicit environmental variable. Application of rule i, above, to all three sources of variance gives 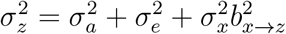
- *σ*_*zw*_: The covariance between the trait and fitness can be decomposed in several ways. The expression given in table 1 presents *σ*_*zw*_ in two components. The first component is the portion of the covariance due to the effect of the trait on fitness, which stems from rule i, above, and the expression for the trait variance, immediately above. The second is the component of trait-fitness covariance resulting from the common effect of the environmental variable on each, which draws on rule iii, above. Taken together, this gives the expression in table 1 of 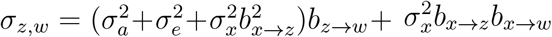.
- *σ*_*x,z*_: The covariance of the environmental variable with the trait follows directly from rule ii, above.
- *σ*_*x,w*_: The covariance of the environmental variable with fitness follows directly from the covariance rule (ii), above, and the total effect rules (2,3) such that the total covariance of *x* and *w* is 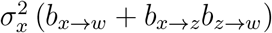.

## Appendix 2: Steps in the demonstration that multiple regression correctly estimates trait-fitness effects when environmental variables are included

Starting with equation 7,

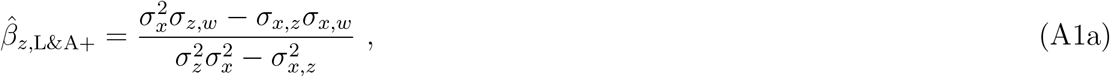

we can show that this estimate of the effect of the trait on relative fitness, accounting for the environmental confounder, coincides with the selection gradient in the quantitative genetic sense (which we have already seen should have a value of *β* = *b*_*z→w*_). Note that 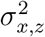 is the square of the covariance of *x* and *z*. First, we substitute expressions for the variance of the trait, and covariances among the trait, environmental variable and fitness, from table 1:

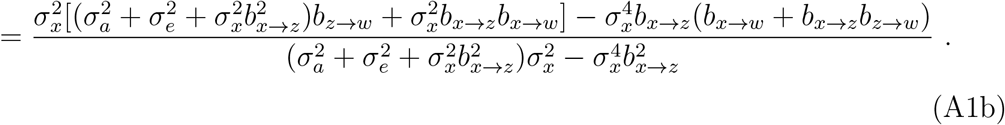

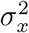 cancels from the numerator and denominator, giving

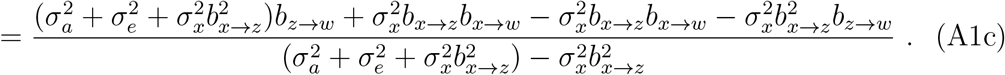

Now, note that 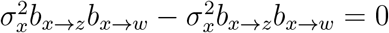, which simplifies the expression to

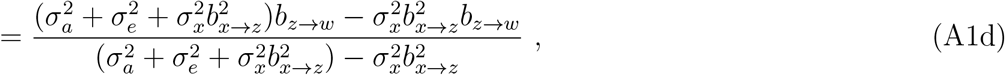

in which we can factor *b*_*z→w*_ from the two remaining terms in the numerator, yelding.

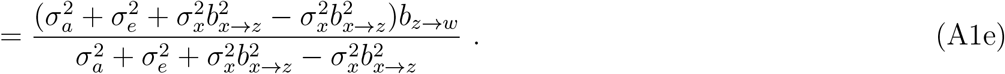

Cancelling like terms in the numerator and denominator yields

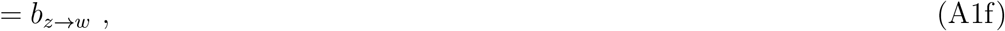

confirming that multiple regression analyses including environmental variables can return estimates of effects of traits on fitness that may be interpretable as selection gradients in the sense that they are valid to use in the Lande equation to predict the course of adaptive evolution.

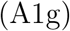

## Appendix 3: Derivation of multivariate (de)compositions of the selection differential in terms of direct and extendedsense selection gradients

The covariance matrix of environmental variables **x** and traits [**u**^*T*^, *f*, **v**^*T*^]^*T*^ by applying equation 14 is

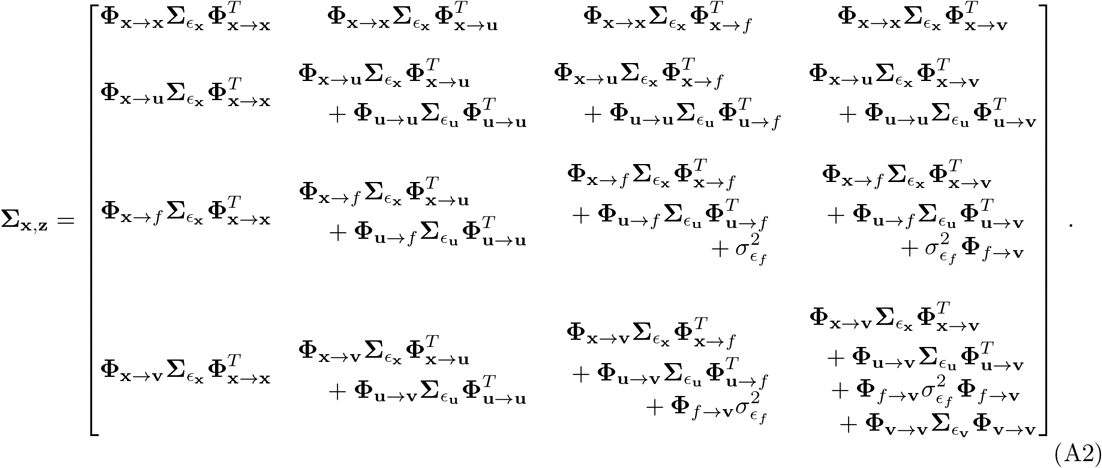

Multiplication of the third row of this matrix by the direct effects of the **x, u**, *f*, and **v** on relative fitness gives equation 15.

The effect of variables causally preceding *f* (i.e., **x** and/or **u**) on traits influenced by *f* (i.e., **v**) can be decomposed into a part acting via 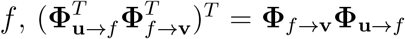 using the effect of **u** as an exmple, and a part not acting via *f*, **Φ**_**u***→***v**_ − **Φ**_*f→***v**_**Φ**_**u***→f*_. Applying this decomposition to the first two terms in part (d) of equation 15 yields

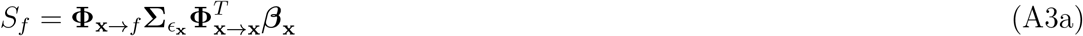

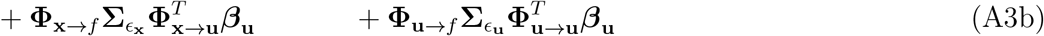

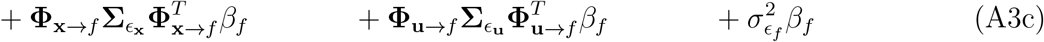

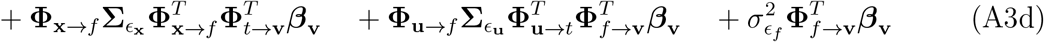

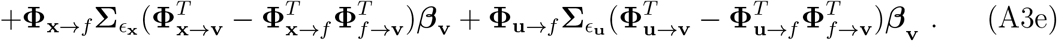

Parts A3a/b describe the part of trait-fitness covariance that arises from from exogenous variance in **x** and **u** jointly influencing the trait and also having direct effects fitness (i.e., those parts of the effects of **x** and **u** on fitness that are neither mediated by *f* nor **v**. Part A3c describes the trait-fitness covariance arising from different sources of variance in the focal trait and the mapping of that trait directly onto fitness via *β*_*f*_ (i.e., not including mediated effects of *f* on *w* via **v**. Part A3d represents all contributions of exogenous variance in **x** and **u** and *f* to covariance of *f*, via fitness effects mediated by both *f* and **v**. Finally, part A3e represents contributions of exogenous variance in **x** and **u** to covariance of *f* and *w*, where the fitness effects are mediated by *f* but not by **v**.

From the decomposition of *S*_*f*_ given in equation A3 rearrangement of lines (c) and (d) yields

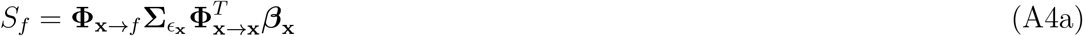

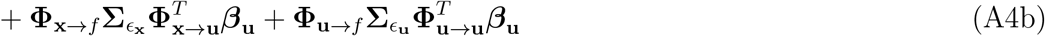

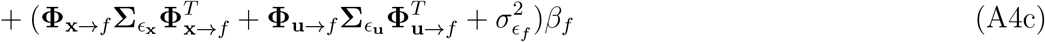

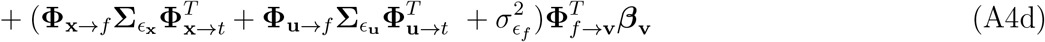

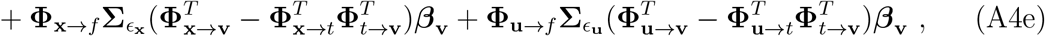

with a common component of parts (c) and (d) that can be factored, noting that

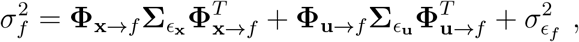

and 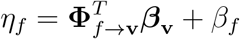, yielding

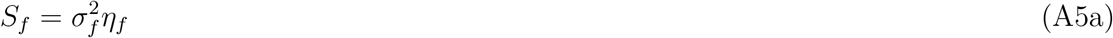

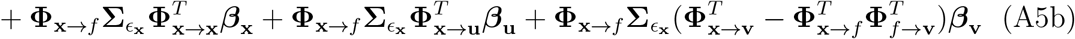

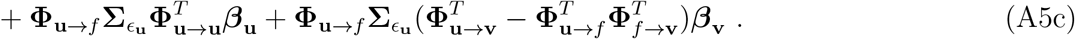

Equation A5 part a represents parts c and d of equation A4, combining the direct effect of the trait (eq. A4c) and the causal component mediated by **b** (eq. A4d) into a total causal effect. However, this also reflects the decomposition of the total causal covariance into direct 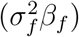 and mediated 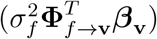 components.

Like terms in equation A5 parts (b) and (c) can be factored,

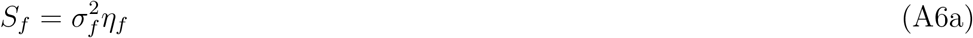

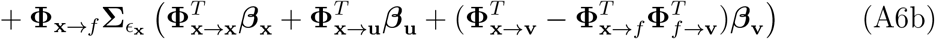

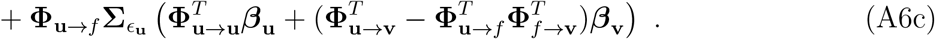

and then adding 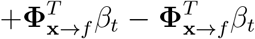 to the terms in large parentheses in part (b) and 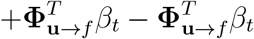 to part (c) gives

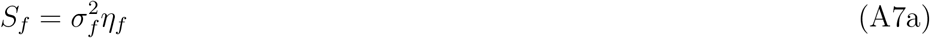

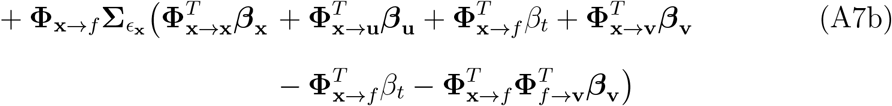

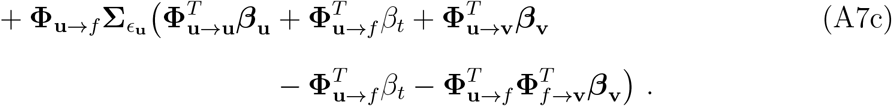

Main text equation 16 is obtained by noting that 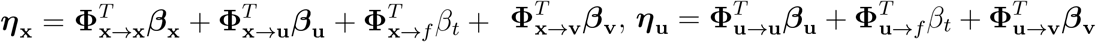, and 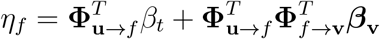.

## Appendix 4: Numerical demonstration of the validity of selection gradient inferences with environmental variables

Variances of exogenous inputs and path effects for the model in figure 1. Variable names are chosen to correspond closely to notation on figure 1:

~~~
> # exogenous variances
> var_a <- 0.5
> var_e <- 0.25
> var_x <- 1
>
> # path effects
> b_zw <- 0
> b_xz <- 0.5
> b_xw <- 0.5
~~~

These values are arbitrary. The following exercises should hold for any values. These specific values illustrate the situation where there is no effect of the trait on fitness, and so all trait-fitness covariance is “evolutionarily inert” environmental confounding.

Simulate a large number of values of the exogenous quantities (*a, e* and *x*), and then the derived quantities (*z* and *w*). Note that the residual variance in *w* is not consequential for any of the quantities that matter for characterising selection and predicting evolution; unit residual variance is simulated to avoid error messages in subsequent application of regression and multiple regression with fitness as the response variable. Note that the intercept of the fitness generating equation is 1. This is necessary because the mean of relative fitness must, by definition, be 1. If the code that generate *z* or *e* was modified generate a trait with a mean other than zero, centred value of *z* would require corresponding modifications to mean-centre *z* and/or *x* before generating the fitness values:

~~~
> # large sample size to converge on long-run behaviour
> n<-10000
>
> # exogenous values
> a <- rnorm(n,0,sqrt(var_a))
> e <- rnorm(n,0,sqrt(var_e))
> x <- rnorm(n,0,sqrt(var_x))
>
> # trait and fitness according to the path model
> z <- a + e + x*b_xz
> w <- 1 + z*b_zw+x*b_xw+rnorm(n)
~~~

Check what evolution would actually obtain, based on the secondary theorem of selection:

~~~
> # the STS
> cov(a,w)
[1] 0.006983459
~~~

The selection gradient estimates from standard and ‘environment-aware’ selection gradient analysis:

~~~
> # standard estimation of beta
> beta_LA <- coef(lm(w∼z))[2]
> beta_LA
Z
0.2557746
>
> # Lande-Arnold-plus estimation of beta
> beta_LAplus <- coef(lm(w∼z+x))[2]
> beta_LAplus
Z
0.007736603
~~~

Evolutionary prediction with the Lande equation, using the `beta_LAplus` selection gradient estimate, should agree well with the secondary theorem of selection. The Lande equation using the `beta_LA` selection gradient estimate may or may not agree, depending on specific values used in the first code snippet in the numerical example

~~~
> round(cov(a,w),3)
[1] 0.007
> round(var_a*beta_LA,3)
z
0.128
> round(var_a*beta_LAplus,3) z
0.004
~~~

## Appendix 5: Decomposition of selection differential in the presence of non-linear effects

Equation 16 decomposes the selection differential into three components, representing (1) the causal effect of the focal trait on fitness, (2) indirect selection due to confounding by other traits that affect both the focal trait and fitness, and (3) indirect selection due to environmental confounding. Here we present a more general version of this decomposition that allows for arbitrary non-linear and non-additive effects of traits and environments. We assume the qualitative structure of causal relationships among environmental variables **x**, traits **u**, *f* and **v**, and fitness *w* that is given in figure 2. The only other substantive assumption is that the focal trait *f* is normally distributed.

In practice, a normal distribution of *f* suggests that **x** and **u** should also be normally distributed and their effects on *f* should be linear. While these conditions are not mathematically necessary for *f* to be normal, normality of *f* would otherwise need particular coincidences of the distributions of **x** and **u** and the functional form of their effects on *f* that do not seem reasonable to presume on biological grounds. Nonetheless, the following derivation allows for all effects to be non-linear and non-additive, so long as *f* remains normal.

For a normally distributed focal trait *f*, we can express the selection differential *S*_*f*_ in terms of the average slope of the fitness curve for *f* (Stein 1981, Walsh & Morrissey 2019):

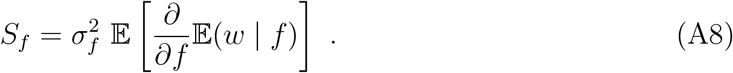

By the law of total expectation, the univariate fitness curve E(*w* | *f*) can be expressed as the integral of the multivariate fitness surface E(*w* | **x, u**, *f*) over the joint conditional distribution of **x** and **u** given *f* :

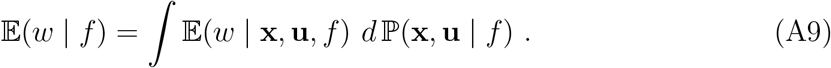

Substituting this expression into equation A8, we can differentiate under the integral sign and apply the product rule of calculus to obtain

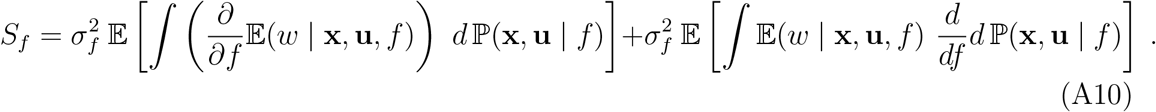

Integrating under the derivative in the first term then yields:

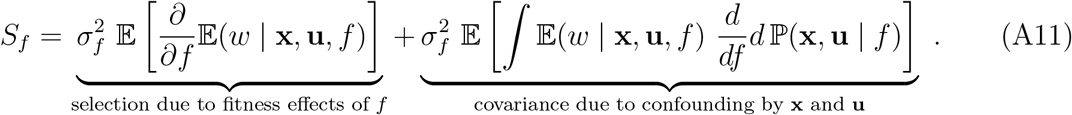

The first term tells us how small changes in the focal trait affect fitness, while controlling for **x** and **u**. Controlling for these variables is sufficient to isolate the causal effect of the focal trait on fitness (i.e., **x** ∪ **u** is a backdoor set for this effect: Pearl 2009, Henshaw *et al*. 2020). This term thus represents the causal component of the selection differential and can be written (Morrissey 2014, Henshaw *et al*. 2020):

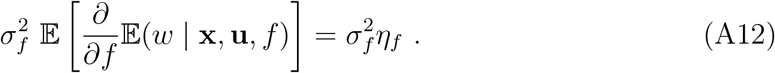

On the other hand, the second term in equation A11 tells us how fitness is affected by changes in the conditional probability distribution of **x** and **u** associated with small changes in *f*. Note that the derivative in the second term should be interpreted as

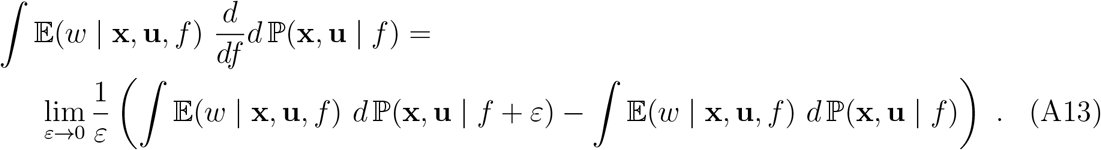

The first term on the right-hand side represents an individual’s expected fitness if their focal trait takes the value *f*, but the confounding variables **x** and **u** are distributed as if the focal trait had taken the value *f* + *ε*. The second term represents the individual’s natural (i.e., non-intervened) expected fitness when the focal trait has value *f*. Taken together, this expression represents the marginal fitness effect of changes in the conditional distributions of **x** and **u** associated with small changes in the focal trait, while holding the actual value of the focal trait fixed at *f*.

Equation A11 decomposes the selection differential into terms representing (1) the causal effects of the focal trait on fitness, and (2) associations due to underlying shared causes, whether these be traits of environmental variables. We can now go a step further to separate indirect selection via other traits from selection due to environmental confounding, as in equation 16. First, by the law of total probability, we have

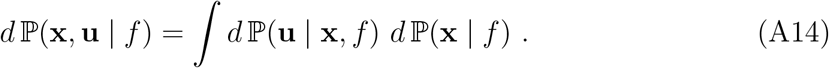

Substituting this expression into the second term of equation A11, we can differentiate under the integral sign and apply the product rule of calculus to obtain:

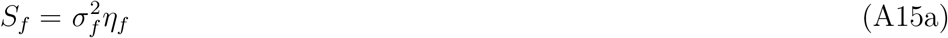

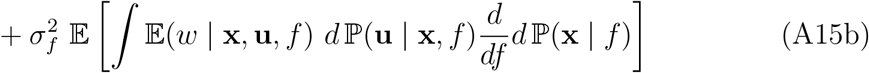

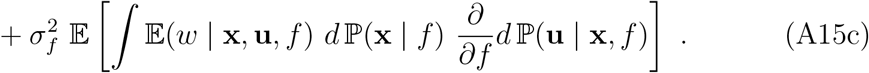

This decomposition of the selection differential is directly analogous to equation 16. Selection on the focal trait is expressed in terms of the total causal effect of the focal trait on fitness (A15a), indirect selection due to confounding by other traits that affect both the focal trait and fitness (A15b), and indirect selection due to environmental confounding (A15c). Importantly, indirect selection may also arise when the focal trait affects other traits (here denoted **v**), which in turn influence fitness. Such effects are included in the total causal effect in equations 16a and A15a. In contrast, the classical framework of Lande and Arnold (1983) does not distinguish between indirect causal effects and confounding by traits: both are considered to result in ‘indirect selection’. Using the above framework, we can further separate the total causal effect of the focal trait on fitness into components acting directly and via **v** (Henshaw *et al*. 2020):

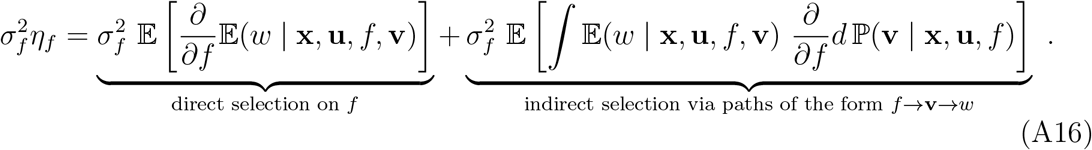

In the notation of Henshaw *et al*. (2020), these terms are given by 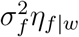 and 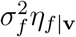 respectively. Note that all of the above quantities are measurable at least in principle, given sufficiently high-quality data. The conditional expectations and probabilities can be estimated using sufficiently general regression techniques (Henshaw *et al*. 2020).

